# Mechanochemical modelling of dorsal closure reveals emergent cell behaviour in tissue morphogenesis

**DOI:** 10.1101/2020.01.20.912725

**Authors:** Francesco Atzeni, Laurynas Pasakarnis, Gabriella Mosca, Richard S. Smith, Christof M. Aegerter, Damian Brunner

## Abstract

Tissue morphogenesis integrates cell type-specific biochemistry and architecture, cellular force generation and mechanisms coordinating forces amongst neighbouring cells and tissues. We use finite element-based modelling to explore the interconnections at these multiple biological scales in embryonic dorsal closure, where pulsed actomyosin contractility in adjacent Amnioserosa (AS) cells powers the closure of an epidermis opening. Based on our *in vivo* observations, the model implements F-actin nucleation periodicity that is independent of MyoII activity. Our model reveals conditions, where depleting MyoII activity nevertheless indirectly affects oscillatory F-actin behaviour, without the need for biochemical feedback. In addition, it questions the previously proposed role of Dpp-mediated regulation of the patterned actomyosin dynamics in the AS tissue, suggesting them to be emergent. Tissue-specific Dpp interference supports the model’s prediction. The model further predicts that the mechanical properties of the surrounding epidermis tissue feed back on the shaping and patterning of the AS tissue. Finally, our model’s parameter space reproduces mutant phenotypes and provides predictions for their underlying cause. Our modelling approach thus reveals several unappreciated mechanistic properties of tissue morphogenesis.

## Introduction

The morphogenesis of tissues and organs is the result of a complex interplay of mechanical and biochemical signalling processes. These control the growth and differentiation states of individual cells in space and time and their cooperative or collective behaviour. A key task is to generate cellular forces that change cell and tissue shape. These forces need to be precisely timed and mechanically balanced in order to generate a desired morphology. In addition, developing tissues are mostly embedded in a tissue environment that itself changes shape with cells following their own morphogenetic programs, which is prone to generate mutual interference. In particular, cells at tissue-tissue boundaries need to adapt to dynamically changing biochemical and mechanical environments and cell-cell contacts. When observing changes in cell behaviour and architecture it is often difficult to separate effects caused by cell autonomous processes and effects superimposed by the environment. Similarly, it is difficult to determine which aspects of cell and tissue behaviour are regulated and which are emergent. Explorative mathematical modelling provides a powerful means to address such intertwined processes.

Dorsal closure (DC) is a relatively simple morphogenetic process occurring in *Drosophila melanogaster* embryogenesis, where modelling has been extensively used in various studies (Kiehart *et al*, 2017; Aristotelous *et al*, 2018). DC comprises the closure of a large opening in the dorsal embryonic epidermis (ES). It can be selectively manipulated and imaged live with sub-cellular resolution, which makes it particularly well suited to address the impact of cellular scale biochemistry and force generation on tissue scale morphogenesis and tissue-tissue interactions. Several force producing mechanisms were identified that occur in the two relevant tissues, the closing ES tissue and the contracting amnioserosa (AS) tissue, which fills the opening (Kiehart *et al*, 2000; Solon *et al*, 2009; Sokolow *et al*, 2012; Saias *et al*, 2015). Selective MyoII activity interference in the two tissues showed that apical constriction of the AS cells provides the critical force for convergence of the epidermis fronts (Pasakarnis *et al*, 2016). When these fronts have come close enough for the dorsal-most epidermis cells (DMEs) to engage, the latter start pulling on each other. This force drives the sequential sealing of the opening, starting at the anterior and posterior canthi from where it “zips” towards the middle of the opening.

Suppression of this zipping force reveals a maximal, autonomous contractility of the AS tissue. The underlying apical constriction process lasts for 3-4 hours and is driven by superimposed periodic apical cell surface area contractions that occur with a periodicity of 2-3 minutes (Solon *et al*, 2009). Dynamic, contractile actomyosin networks transiently forming at random positions of the apical plasma membrane produce this oscillatory behaviour (Blanchard *et al*, 2010). It is not clear how these AS cell surface area oscillations begin translating into apical constriction at the onset of closure.

Coinciding with this transition is the beginning of an oscillation amplitude decrease, which continues until cells eventually arrest. This occurs in a patterned manner, with the smallest AS cells at the anterior and posterior canthi of the eye shaped opening arresting first, followed by the ventral-most cells and then sequentially, by the more centrally situated (Solon *et al*, 2009; Blanchard *et al*, 2010). A gradient of dpp, secreted by the dorsal-most ES cells comprising the boundary to the AS tissue (DMEs), was proposed to control apical constriction of AS cells (Lada *et al*, 2012). In addition, mechano-chemical coupling of neighbouring AS cells was suggested to provide tissue autonomous regulation (Hunter *et al*, 2014). Furthermore, genetic interference with actomyosin dynamics, suggested additional, cell autonomous regulation, where the periodic bursts of apical F-actin formation in AS cells require a feedback from contractile non-muscle myosinII (MyoII) activity (Azevedo *et al*, 2011; David *et al*, 2013; Saias *et al*, 2015).

Here, we question such a role for MyoII, as we show experimentally that oscillatory F-actin dynamics occur in the absence of MyoII activity and mechanical coupling. Based on these findings, we established a multi-scale modelling approach that, unlike previous modelling of DC, integrates biochemistry governing subcellular scale actomyosin organisation and the mechanics of MyoII-driven AS cell- and tissue shape changes (Aristotelous *et al*, 2018). We implemented our model using the finite element method (FEM), which can handle complex geometries and their deformation through discretisation into numerous, simple elements (e.g. triangles, quadrilaterals, tetrahedra). FEM-based simulations have been used to describe how complex biological structures form during development (Brodland & Clausi, 1995; Davidson *et al*, 1995). Recently, FEM was also used to simulate reaction diffusion equations coupled to material contractility on simple spherical shapes (Brinkmann *et al*, 2018). We here similarly combined tissue mechanics with cellular actomyosin biochemistry in AS cells, segmented from a living embryo, to simulate and explore AS tissue dynamics during DC.

Exploring the model’s free parameter space revealed that depleting MyoII activity can indirectly affect F-actin dynamics without the need for direct feedback on F-actin biochemistry, as generally assumed. Furthermore, it predicted emergence of a previously described, graded AS cell pulsation arrest, which so far seemed controlled by a corresponding Dpp signalling gradient. Supporting the model’s prediction, AS tissue-specific interference with Dpp signalling *in vivo*, showed no effect on DC. In addition, the model showed that both, the pattern with which AS cells arrest and the overall AS tissue shape changes critically depend on the mechanical properties of the surrounding epidermis tissue. Finally, several of the thousands of solutions, provided a clear cause for some of the cell and tissue phenotypes that had been reported for genetic mutants. They thus make a specific prediction for the role of the respective proteins. Altogether, our modelling approach generates several unexpected mechanistic views on morphogenesis in general and DC in particular.

## Results

### Periodic F-actin dynamics are MyoII-independent

The oscillating actomyosin dynamics of AS cells were proposed to depend on MyoII activity but so far supporting experimental data have not been shown (David *et al*, 2013). We therefore tested this by selectively interfering with MyoII activity using the deGradFP system in AS cells (Caussinus *et al*, 2012; Pasakarnis *et al*, 2016). These expressed the mCherry tagged F-actin-binding domain of Moesin to visualise F-actin dynamics (mCherry-moesin). As previously shown, the AS cells of such AS-SqhKO embryos completely lacked MyoII dynamics, showed no apical cell surface oscillations and could not apically constrict thus failing DC (Pasakarnis *et al*, 2016). Surprisingly, we now found that periodic F-actin dynamics were not much affected in AS-SqhKO embryos (Fig. 1A; Movie 1; Methods). Similar to the wild type, local F-actin foci formed in all cells and migrated across the apical cell surfaces similar to the wild type, before vanishing again. Generally, the appearing F-actin foci had reduced signal intensity. However, this is expected, as due to the lack of MyoII activity these foci will not contract. In the wild type, such contraction concentrates the actin filaments, which locally enhances the detectable signal. Compared to the wild type, the periodicity of F-actin foci formation was only slightly longer (Fig. 1B). This result contrasts a previous study mentioning the absence of periodic F-actin nucleation in embryos that were homozygous mutant for *zipper* (*zip*^*1*^), the gene encoding MyoII heavy chain (David *et al*, 2013). However, when imaging *zip*^*1*^ embryos expressing mCherry-moesin, we found periodic F-actin dynamics very similar to those in AS-SqhKO embryos (Fig. 1C). These data show that the periodically forming F-actin networks of AS cells are not dependent on MyoII activity.

**Figure 1:**
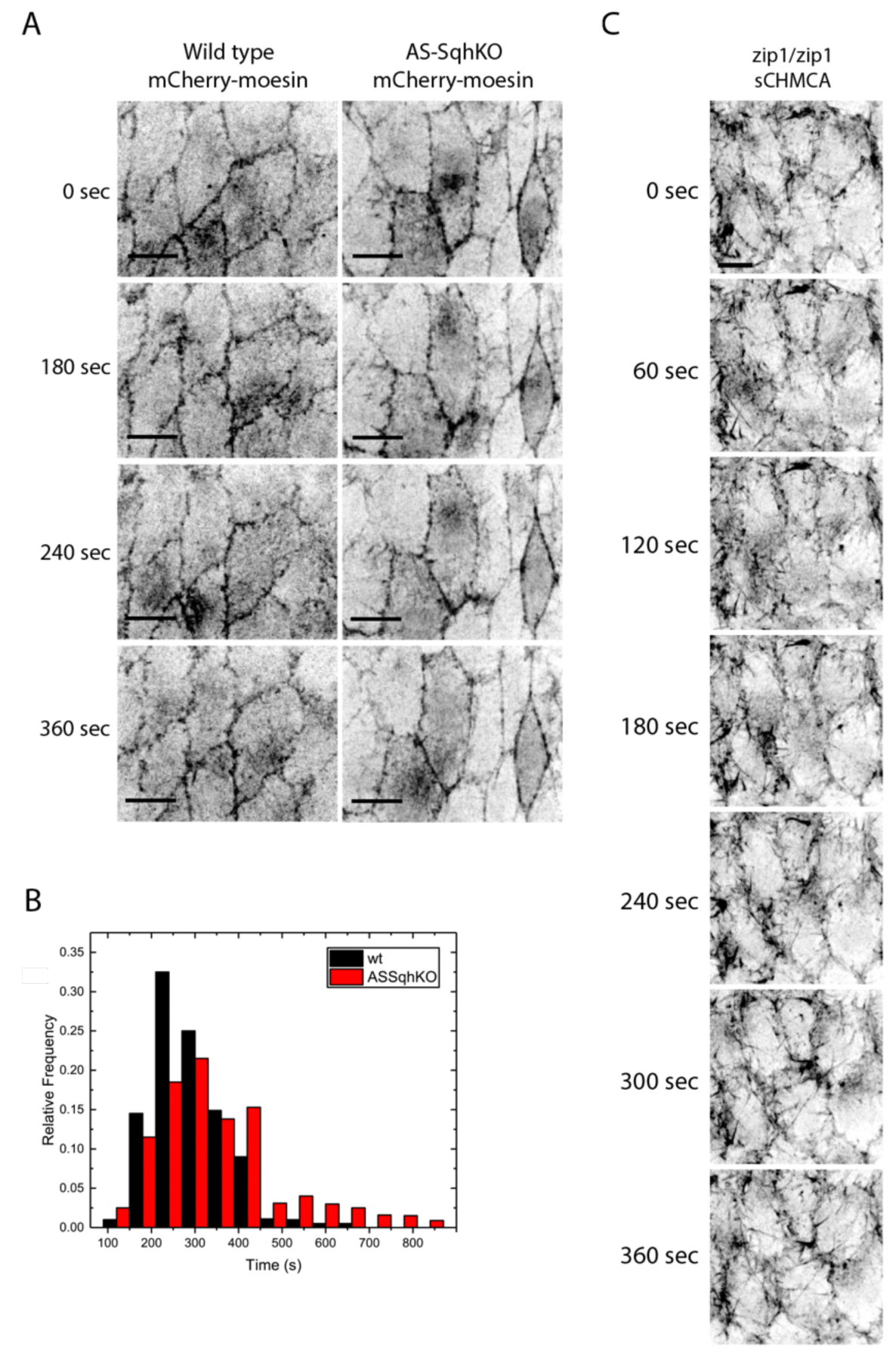
F-actin oscillation are Myosin-independent. A) Comparison of F-actin dynamics wild type (upper row) and MyoII activity-depleted amnioserosa cells (lower row) in AS-SqhKO embryo expressing mCherry-moesin to visualize F-actin. Scale bars: 20 µm. B) Quantification of F-actin dynamics comparing wild type and MyoII activity-depleted amnioserosa cells of AS-SqhKO embryos. C) F-actin oscillations in AS cells homozygous mutant for *zip1* encoding the MyoII heavy chain. Scale bars: 10 µm.

### FE modelling of F-Actin dynamics in AS cells

A crucial question to address is how the relatively slow apical constriction connects to the much faster oscillating actomyosin contractility. To date, experimental limitations have hampered an unambiguous answer. For this reason, we decided to develop a computer-based, mathematical modelling approach that would allow building a tissue system to explore the mechanistic link between actomyosin activity, cellular force production, force transduction and tissue morphogenesis. For this, we combined biochemical reactions relevant to cytoskeleton dynamics with cell and tissue mechanics in a FEM-based implementation of the DC process. In our custom program, we implemented an image-based discretisation of roughly 150 cells comprising a typical AS tissue. The surrounding ES tissue was similarly discretised, but without subdividing it into cells (Fig. 2A). As a basis, we generated a region that was entirely subdivided into linear hyper-elastic triangular elements on which the mechanical problem was solved using the finite element method (Appendix Supplementary Methods). The region was further subdivided such that an outer ring of “mechanical elements” constituted the ES tissue while the central area was subdivided into cells constituting the AS tissue. The mechanical elements of each AS cell were further subdivided, to generate a finer “biochemical” mesh, onto which we implemented the actomyosin activity (Fig. 2A). Actomyosin modelling was exclusively performed in the AS tissue, while the contribution of the surrounding ES tissue was purely mechanical and thus passive. We modelled apical cell surfaces only, where the relevant actomyosin dynamics and force transmission are known to take place (Martin & Goldstein, 2014). The biochemistry was implemented with a reaction-diffusion equation-based model that produced the periodic formation of F-actin foci in a spatially resolved manner (Appendix Supplementary Methods). Diffusion of the components was restricted by cell boundaries. Based on our *in vivo* experiments, we designed a model for oscillating F-actin formation dynamics that is independent of MyoII. We simplified F-actin nucleation by integrating the activity of F-actin nucleators and the respective regulators into a species we term Actin Nucleating Proteins (ANPs). These are produced in an inactive state (ANPi) and can become activated (ANPa) (Fig. 2B). To model spontaneous, local triggering of ANP activation, we use a Turing patterning mechanism (Turing, 1952) (Appendix Supplementary Methods). The reactions were built on the activator-depleted substrate model from Gierer and Meinhardt, 1972 (Gierer & Meinhardt, 1972). These ANP dynamics were then coupled to an additional species that modelled F-actin.

**Figure 2:**
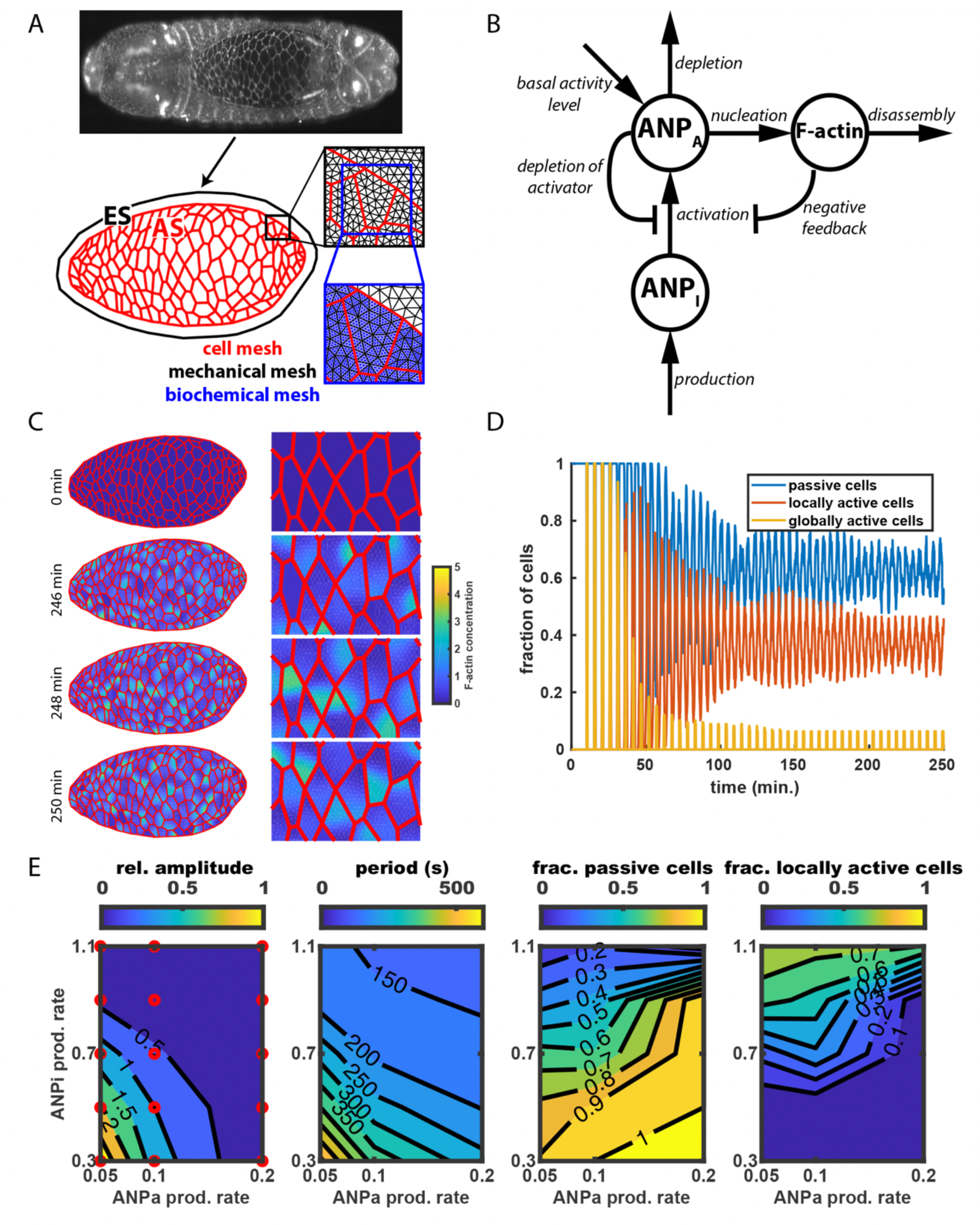
Modelling Myosin-independent F-actin oscillation with reaction-diffusion equations. A) The mesh generation procedure: AS cells were manually segmented starting from the maximum projection of a SPIM image and a strip of epidermis (ES) was arbitrarily defined around the AS; the mechanical mesh was computed for both AS and ES tissues; the biochemical mesh was computed as a subdivision of the mechanical mesh for AS only. B) Reaction diagram for F-actin dynamics in the model. Pointed arrows indicate positive feedback, constant or linear reaction. Capped arrows indicate negative feedback. C) F-actin conc. at multiple time steps in absence of myosin activity. D) Fraction of cells in each F-actin activity category over time. E) parameter screens statistics, left to right: relative amplitude and period of cells’ F-actin oscillation; passive and locally active fraction of cells at equilibrium. Each plot shows average values over all AS cells and time steps, with contour lines calculated with MATLAB’s contour function, starting from measures of simulation runs (red circles).

Our model comprised the following interactions, the equations of which we provide in the Appendix Supplementary Methods: ANPi is produced at a constant rate and is activated to become ANPa at a rate that depends on ANPi and ANPa concentration (Fig. 2B). ANPa leads to F-actin formation at a linear rate. Similarly, ANPa depletion and F-actin disassembly occur at a linear rate. Furthermore, we assume negative Hill-type F-actin feedback on ANP activation. This introduces the previously reported negative feedback of F-Actin on the activity of Rho-GTPases (Bement *et al*, 2015; Michaux *et al*, 2018; Segal *et al*, 2018). Notably, in our simulations we found this feedback to increase the number of parameter combinations that produce oscillatory dynamics suggesting that this feedback merely serves to provide additional robustness (Appendix Supplementary Methods). Consistent with the original Gierer-Meinhardt model, we implement a negative Hill-type feedback on ANP activation, which eventually leads to a saturation of ANPa production. ANPa concentration cannot drop below a minimal level, which not only improved the stability of computer simulations but also provided a parameter, the modulation of which allowed simulating certain mutant *in vivo* scenarios (see below).

Having found that actin bursts and waves were present in the absence of MyoII *in vivo*, we first explored the parameter space of our reaction-diffusion model to identify the conditions that qualitatively reproduced a similar oscillatory F-actin behaviour. To home in on the greater area of the parameter space of interest, we first fixed the parameters of the reaction part of the equations using standard dynamical systems theory (Murray, 2004). Next, we fixed the ANP diffusion coefficients. For the ANPi diffusion coefficient, several publications suggest that 10µm^2^/s is a reasonable value (Mogilner & Edelstein-Keshet, 2002; Holmes *et al*, 2012). The ANPa diffusion coefficient should be considerably lower, as much of the ANPa will be F-actin bound. Lacking sufficient relevant experimental information, we again used theoretical methods to calculate the range of ANPa diffusion coefficients that would lead to oscillatory F-actin dynamics as observed *in vivo* (Appendix Supplementary Methods).

We applied this system of reaction-diffusion equations to the tissue geometry that we had extracted from an image of a living embryo (Fig. 2A). For this, we chose the time point of DC when the two lateral ES tissue fronts had just made their initial contact at the anterior end of the opening to fully enclose the AS tissue. Simulations were initialised with starting concentrations of ANPi, ANPa and F-actin set to zero (Fig. 2C). To quantify the subcellular spatial pattern of F-actin over time, we extracted the cell area fraction that was covered by F-actin at each time point: Below 20% coverage we considered a cell to be passive, above 60% coverage to be globally active and in between to be locally active (Appendix Supplementary Methods). Initially, all cells simultaneously increased their ANPi, and consequently their ANPa and F-actin concentration, causing synchrony of the first F-actin oscillation cycle (Fig. 2D). Cells gradually de-synchronised in the following cycles until oscillations reached an equilibrium distribution (Fig. 2D; Movie 2). Thereby, cells alternated between passive and locally or globally active states over time (Fig. 2C). Once the initial synchrony of activation was broken, the average fraction of cells in each of the categories converged to a stable value at any given time point (Fig. 2D). To explore the dependence of this equilibrium value on the chosen parameters, we systematically changed the production rates of ANPi and ANPa (Fig. 2E). The resulting 2-dimensional parameter spaces showed that for low values of both parameters most cells were passive. The other cells had high oscillation amplitudes and periods. As the ANPi production rate increased, the number of passive cells gradually decreased, while the number of cells with F-actin oscillations increased (Fig. 2E). Concomitantly, these cells showed decreasing periodicity and amplitudes. In summary, our conceptualisation of a basic F-actin biochemistry, for defined parameter values, is able to produce oscillatory F-actin dynamics that resemble the dynamics observed *in vivo*, in wild type and MyoII mutant AS cells.

### MyoII forces modulate oscillatory F-actin dynamics

After our modelling system reproduced F-actin oscillations, we went on to explore how these oscillations react to MyoII force action, by connecting the biochemical and mechanical finite element models (Fig. 2A). Since oscillatory F-actin dynamics occur independent of MyoII activity, there is in principle no need for additional MyoII regulation. Therefore, we assumed that the motor protein by default binds to F-actin and is present in an active form at non-limiting concentration. In this scenario, the oscillating F-actin concentration is the critical variable controlling contractility (Fig. 3A). In addition, we assumed that we need a critical amount of F-actin bound MyoII, in order to produce a force that is sufficient to initiate contraction. Such a relationship between F-actin and contractility can be described with a Hill function, whereby contractility sharply increases until a maximal value (cMax) once F-actin concentration is above a threshold concentration (cFthr) (Appendix Supplementary Methods). In our finite element implementation this means that the overall F-actin concentration of all biochemical finite elements within a mechanical finite element governs its contractility. Notably, the degree of contraction of a mechanical element in addition is governed by the elastic material properties and the contractility of the surrounding elements (Appendix Supplementary Methods). We assumed that any cell surface area changes introduced by MyoII activity will not affect the amount of ANPi production, which remained constant throughout the closure process. *In vivo*, this would reflect that the amount of F-actin nucleators reaching the apical cell surface is constant, independent of surface size.

**Figure 3:**
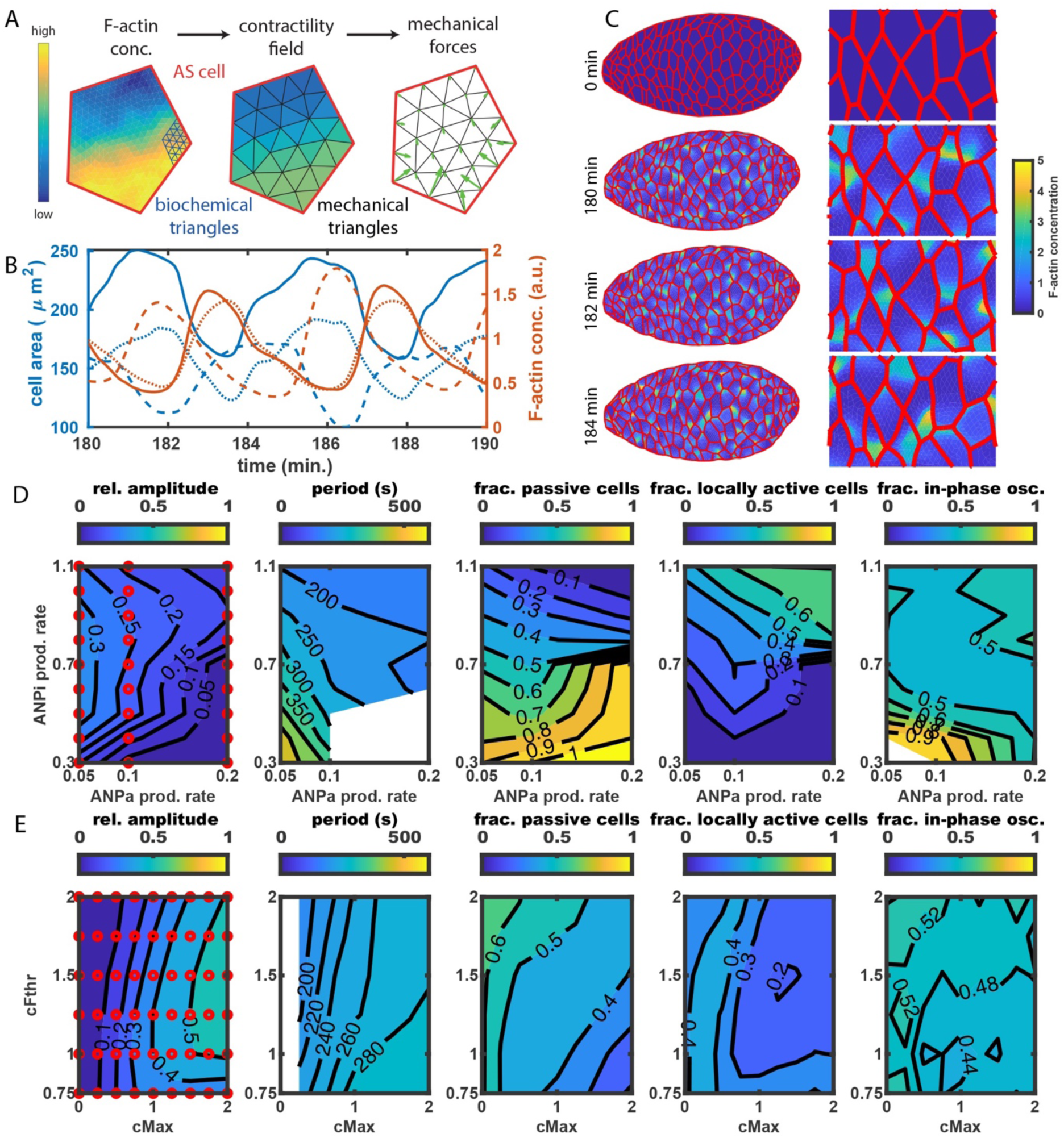
Simulations provide interpretation for WT and perturbed in vivo phenotypes. A) Schematic representation of contractility calculation: first, F-actin conc. is calculated on the biochemical mesh; then, the concentration is extrapolated to the larger mechanical triangles; last, mechanical forces and corresponding deformation are calculated. B) Cell area and F-actin over time for 3 cells at the center of the opening. C) F-actin conc. at multiple time steps with myosin activity. D, E) parameter screens statistics, when varying production rates of ANPa and ANPi (D), and the contractility parameters cMax and cFthr (E). Statistics plot are, left to right: relative amplitude and period of cell area oscillation; inactive and locally active fraction of cells at equilibrium; fraction of in-phase neighbor oscillations. Each plot shows average values over all AS cells and time steps, with contour lines calculated with MATLAB’s contour function, starting from measures of simulation runs (red circles).

We first ran simulations implementing the ES tissue surrounding the AS tissue as a linear elastic material. In this way, the ES tissue provided resistance to the closure forces generated by the AS cells. In our model, changes of F-actin concentration instantaneously translated into contractility changes, such that cellular F-actin concentration peaks and cell area minima occurred simultaneously, and *vice versa* (Fig. 3B). Again, cellular F-actin oscillations started synchronously but rapidly converged towards a non-synchronous equilibrium (Fig. 3C, Movie 3). Interestingly, for a given set of reaction diffusion equation parameters, the added contractility changed the equilibrium features. In particular, we found that certain combinations of ANPi and ANPa production rates produced oscillating F-actin activity only after contractility was added (Fig. 3D). Hence, our model predicts that also *in vivo*, conditions can exist under which MyoII activity becomes essential for periodic F-actin nucleation even in the complete absence of any biochemical feedback (Fig. 2E and 3D).

As in the simulations lacking contractility, increasing ANPi and ANPa production rates in the presence of contractility decreased the average relative amplitude and the average period of oscillations, albeit not to a similar extent (Fig. 2E and 3D). In contrast however, the added contractility influenced the fraction of cells with local or global F-actin activity. For ANPi and ANPa production rates larger than those included in Fig. 3D, less and less cells were passive, and gradually, all of them became globally active. While it seems trivial that with increasing ANPi and ANPa production rates, also the frequency of F-actin structures in cells increases, it is less obvious that this results in oscillations with smaller amplitudes. Notably, this again shows how MyoII activity can indirectly modulate cellular F-actin behaviour without direct biochemical feedback.

The emergent influence of MyoII activity on F-actin dynamics made us address how the latter depends on the key parameters determining contractility, cMax and cFthr. Increasing cMax, which represents increasing MyoII activity, progressively decreased the fraction of passive cells, independent of cFthr (Fig. 3E). This is caused by a combination of two effects: First, the contraction process locally increased the F-actin concentration and second, F-actin structures became more persistent, with the reaction diffusion equation parameters modulating this latter effect. Furthermore, we found that with increasing cMax, also the average oscillation period increased, at least for relatively high values of cFthr. This indicates that the effect of persistent F-actin structures dominates the contraction-mediated effect. Notably, the lowest tested values of cFthr biologically do not make much sense as they would correspond to MyoII-driven contractility at low F-actin concentration. As cFthr was increased from such low values to higher values, the average amplitude of oscillations rapidly increased, while the fraction of passive cells decreased (Fig. 3E). This occurs because high cFthr corresponds to MyoII-binding only taking place in regions with sufficiently high F-actin concentrations, which results in strong local contractions that can deform neighbouring cells. This scenario reflects the *in vivo* behaviour of AS cells (Solon *et al*, 2009; Blanchard *et al*, 2010).

Perturbing actomyosin dynamics in simulations, enabled us to address the coordination of actomyosin dynamics between neighbouring cells. Experimental investigation had shown that neighbouring cell areas mainly oscillate either in anti-phase or in-phase, with a preference for the former (Solon *et al*, 2009). Later, the orientation of neighbouring cells was suggested to determine the preferred mode of coordination, causing stripes of cells to contract in phase (Blanchard *et al*, 2010). In our modelled system, the sub-cellular actomyosin dynamics of a given cell and the resulting contractility, influence the actomyosin dynamics of neighbouring cells by changing their geometry independent of chemical cell-cell coupling. We wondered whether such mechanical coupling between cells would lead to any preferred phase shift between neighbouring cells. To quantify such phase shifts, we used the amount of F-actin close to the junction between the two cells as a proxy for contractility (Appendix Supplementary Methods). In-phase oscillations corresponded to F-actin concentration peaking at the same time in neighbouring cells, while anti-phase oscillations corresponded to a peak occurring in one cell while the neighbour was in a valley. Varying the production rates of ANPi and ANPa in simulations with MyoII activity, revealed a prevalence of in-phase oscillations in correlation with higher amplitudes (Fig. 3D). However, if contracting regions covered larger subcellular areas, we observed a tendency towards anti-phase oscillations (Fig. 3D). In contrast, varying cMax and cFthr did not produce any clear trends neither concerning oscillation amplitudes and period nor fraction of passive and locally active cells (Fig. 3E). In summary, we find that variation of few parameters in our system generated a wide variation of oscillation coordination patterns between neighbouring cells, simply due to added contractility and without the need for biochemical coupling.

### ES tissue relaxation provides DC progression and emergent AS cell pulsation arrest

The previous simulations treat the ES tissue as a passive, elastic material that provides increasing resistance to the closure forces generated by the AS cells. Consequently, an equilibrium resulted, which rather quickly halted closure. *In vivo* however, several rows of ES cells at the AS tissue boundary elongate significantly along their dorsal-ventral axis, suggesting that the ES tissue relaxes during DC (Riesgo-Escovar & Hafen, 1997). To explore this possibility we implemented gradual ES tissue relaxation in our simulations. Since a time-resolved, rheological characterization of ES cells is missing, we assumed simple Maxwell-like relaxation at a constant relaxation rate (Appendix Supplementary Methods). The relaxation timescales used in this work were chosen such that simulated and *in vivo* closure progressed similarly. These were in the order of tens of minutes, which is only slightly larger than what was experimentally measured in other comparable systems (Étienne *et al*, 2015; Doubrovinski *et al*, 2017; Atzeni *et al*, 2019). We further assumed that relaxation is isotropic everywhere in the ES tissue, except along the anterior-posterior axis (Fig. 4A). The latter accounts for the fact that *in vivo* the AS tissue does not shorten in the anterior-posterior direction during DC (Harden *et al*, 2002). Before activating relaxation, we ran each simulation without relaxation for a time period bringing the system close to the oscillations equilibrium (see previous section). This scenario is realistic as *in vivo*, the AS cells oscillate around a constant opening area prior to the onset of ES cell elongation and tissue convergence (Blanchard *et al*, 2010).

**Figure 4:**
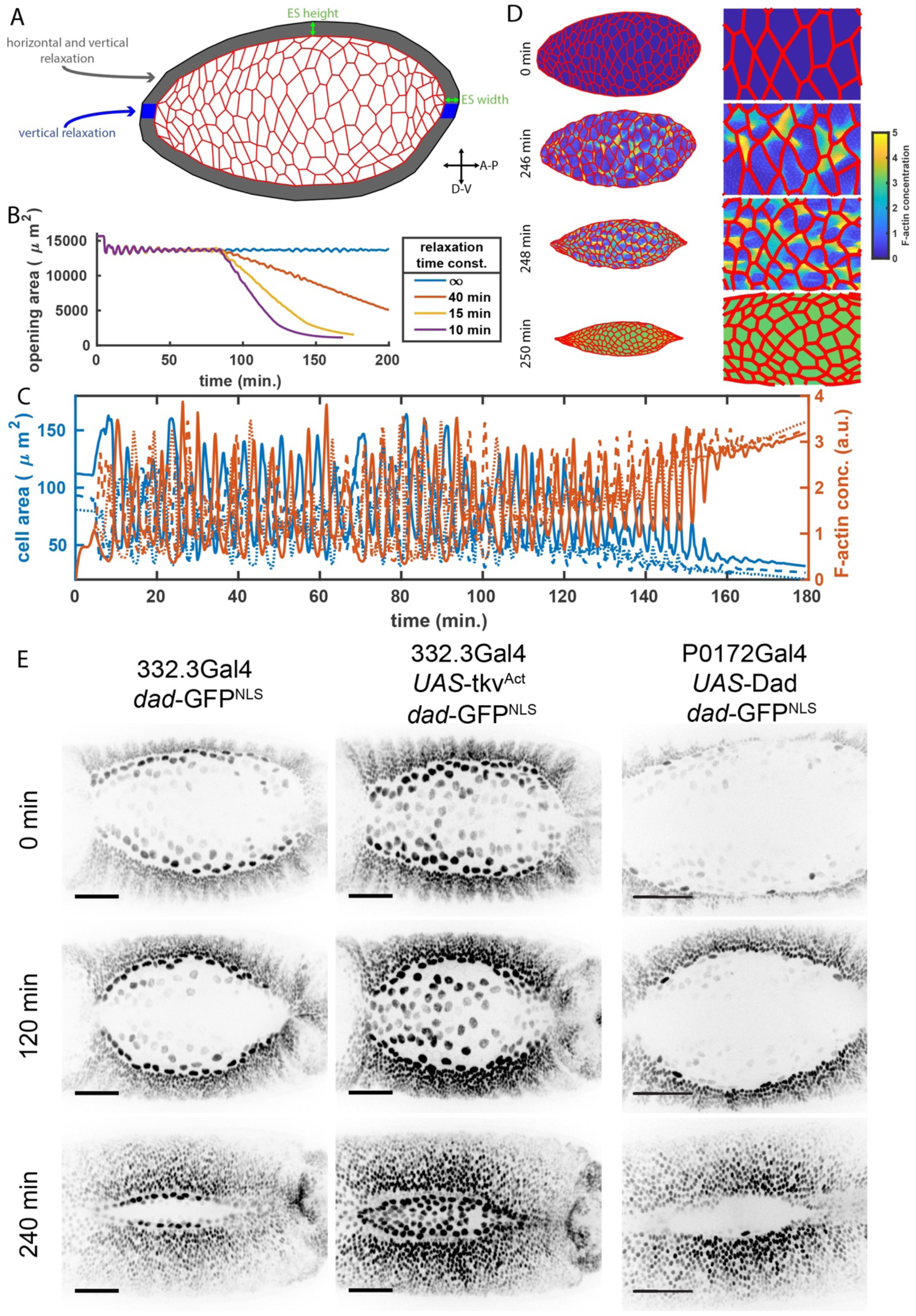
ES Tissue relaxation mediating Dpp-independent AS tissue contraction. A) Choice of ES regions with isotropic and anisotropic material relaxation. B) AS tissue area dynamics with different relaxation time constants. C) Cell area oscillations (blue) in correlation with F-actin concentration dynamics (red) for 3 randomly chosen cells (solid, dashed and dotted lines). D) Sub-cellular F-actin concentration dynamics at indicated time points in a simulation including Myosin activity and ES material relaxation. E) As tissue-specific interference with Dpp signalling gradient by overexpression of constitutively active Dpp receptor thickveins (tkv^act^, second column) or the signalling inhibitor Dad (third column) in embryos expressing nuclear GFP under control of the Dpp responsive dad promotor to visualise cellular Dpp signalling activity levels. In all cases closure occurs similar to the wild type (first column).

Implementing relaxation was sufficient to provide closure progression. In the simulations, the opening area first decreased slightly before stabilising around an equilibrium given by the ES tissue elasticity. When relaxation started the opening area continued decreasing with a rate that was not only set by the ES tissue relaxation rate, but also by the parameters of the reaction-diffusion equations (Fig. 4B, Movie 4). If the relaxation timescale was sufficiently fast, we observed that cell pulsing eventually arrested coinciding with the cessation of F-actin oscillations (Fig. 4C). Thereby, cells transitioned from the dynamic spatial F-actin organisation distribution at the initial equilibrium to a uniform, high F-actin concentration covering their entire surfaces (Fig. 4D, Movie 4). Interestingly, this emergent F-actin oscillation arrest was temporally and spatially patterned: the first cells to reach complete F-actin coverage were generally directly at or very close to the tissue boundary. From there, the behaviour progressed through the tissue towards the middle of the opening as if following a gradient. Intriguingly, a similar sequential pulsation arrest of AS cells had been observed to occur *in vivo*, where it was associated with the maintenance of tissue tension (Solon *et al*, 2009). In our simulations, the consequence of this sequential pulsation arrest was a slowing down of overall AS tissue constriction until closure eventually came to a halt when reaching the maximal effect of contractility. This phenocopies the closure behaviour of embryos in which the force contribution of the zipping process was suppressed (Jankovics & Brunner, 2006).

### Dpp signalling interference in AS tissue does not affect DC

Our simulations predict that graded AS cell pulsation arrest is an emergent property of the system. *In vivo* however, it seemed plausible that such behaviour was controlled by a gradient of the Dpp signalling protein that is known to be secreted at the onset of DC by the DME cells experiencing Jnk signalling (Riesgo-Escovar & Hafen, 1997). Consistently, zygotic mutants of the Dpp receptor thickveins (tkv), amongst other defects, were shown to interfere with AS cell apical constriction but not apical cell surface oscillations (Fernández *et al*, 2007). This was linked to Dpp’s role in contributing to the expression of the MyoII heavy chain component zipper (zip) in AS cells (Zahedi *et al*, 2008). A Dpp signalling-mediated gradient of MyoII protein levels and activity could well account for the sequential arrest of AS cell surface oscillations. To investigate the discrepancy between simulations and *in vivo* experiments, we turned to *in vivo* experimentation. We first monitored DC in embryos expressing GFP, equipped with a nuclear localisation sequence, under the control of a dad enhancer element (*dad*-GFP^NLS^) that selectively responds to Dpp signalling activity (Ninov *et al*, 2010). In wild type embryos, *dad*-GFP^NLS^ revealed a clear Dpp signalling activity gradient across the AS tissue that persisted throughout DC (Fig. 4E; Movie 5). A differential Dpp response already became visible in AS cells with the beginning of germ band retraction, which contradicts previous findings suggesting a first round of Dpp exposure occurring shortly after germ band extension (Movie 5) (Garcia Fernandez *et al*, 2007). Well established is the Dpp signalling event induced by JNK activity in the DMEs at the onset of DC (Riesgo-Escovar & Hafen, 1997; Fernández *et al*, 2007). This second Dpp activity became visible following a short period of constant *dad*-GFP^NLS^ signal intensity. It had limited effect on the AS tissue as only the AS cells closest to the DMEs further increased their nuclear GFP signal in the following hours (Movie 5). In contrast, the nuclear GFP signal in multiple rows of the surrounding ES cells started gradually increasing with DC onset, eventually becoming very bright towards the end of the process. This suggests that a graded response of AS cells to Dpp signalling is established prior to DC, at around the onset of germ band retraction and that the second Dpp signalling activity during DC mainly targets the cells of the ES tissue. Furthermore, it indicates continued Dpp signalling by DMEs throughout DC.

Potentially, the graded exposure of AS cells to the first Dpp signalling event could still determine the apical constriction behaviour of AS cells later on. A problem with previous studies interfering with Dpp signalling was that they used zygotic mutants or global RNAi interference to perturb Dpp activity. Interference thus lacked temporal control and tissue specificity, which hampers the unambiguous interpretation of the resulting phenotypes. To circumvent this problem and specifically explore the requirement of Dpp in AS cells, we selectively interfered with Dpp signalling in the AS tissue (Materials and Methods). First, we ectopically expressed a constitutively active form of the Dpp receptor thickveins (Tkv^Act^) in the AS tissue of embryos expressing *dad*-GFP to monitor Dpp signalling activity. This gradually enhanced *dad*-GFP expression in AS cells during DC, such that also cells in the tissue centre showed a strong nuclear GFP signal (Fig. 4E). Despite the increased Dpp signalling in AS cells, DC proceeded normally. Next, we supressed Dpp signalling by overexpressing Dad, an inhibitor of the Dpp signalling response in the AS tissue (Tsuneizumi K., Nakayama T., Kamoshida Y., Kornberg T.B., Christian J.L., Tabata T. (1997) Daughters against Dpp modulates Dpp organizing activity in Drosophila wing development Nature 389:627-31). This almost completely abolished Dpp signalling response in the AS tissue but again did not affect DC (Fig. 4E). Together these results suggest that Dpp signalling does not critically affect AS tissue behaviour.

### AS and ES tissue properties determine tissue patterning and shaping

Having found good evidence that differential AS cell behaviour is an emergent property of the system, we investigated which parameters and assumptions in our model were critical for the sequential arrest of cell surface oscillations. We defined the time of pulsation arrest as the last time step in which the cell is not fully covered by F-actin. We assumed that ES tissue relaxation and thus DC begins after 80min of simulation time and set a relaxation time constant of 10min to allow closure to proceed within a reasonable computational time (Fig. 4B). Simulations were run for a maximum of a day of computational time or 250 minutes of simulation time. First, we varied the contractility parameters, as before when exploring the role of MyoII. When cFthr was high and cMax low, cell areas oscillated but only very few AS cells showed pulsation arrest (Fig. 5A). The number of arresting cells increased if simulations were run for longer or if faster ES tissue relaxation was implemented (Appendix Supplementary Methods). This occurred also if cMax was increased, for any given value of cFthr. In this scenario, cells arrested sequentially, starting from the outside towards the middle of the opening. Interestingly, if cMax increased further at low cFthr values, some cells stopped oscillating even before the beginning of ES tissue relaxation. When both cFthr and cMax were high, the cell’s oscillation amplitudes were much increased and passive phases were extended (Fig. 3E). This caused faster AS tissue contraction, while area differences between cells were also larger and shorter lived. At the same time, pulsation arrest was more synchronous.

**Figure 5:**
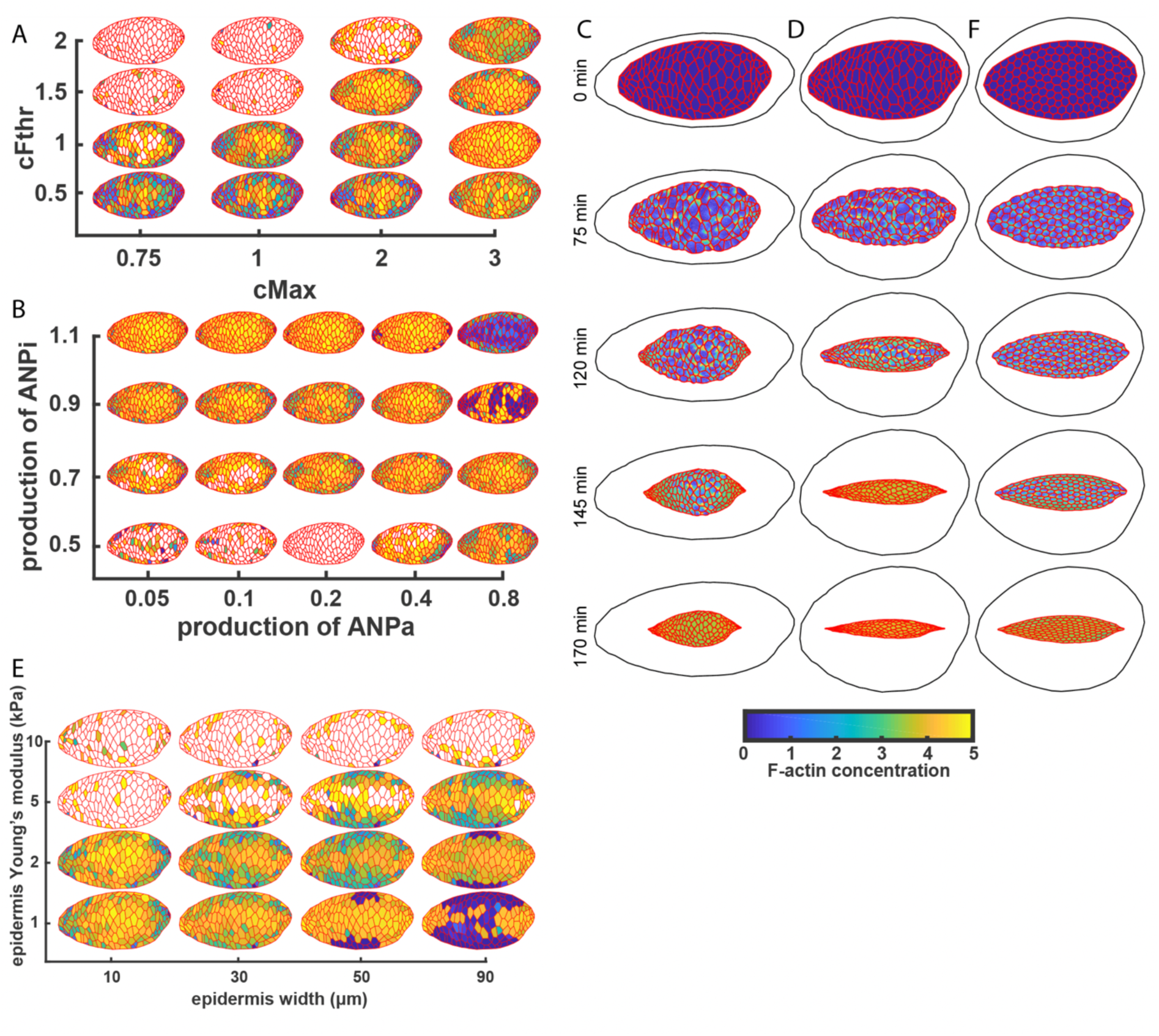
Exploring ES and AS properties reveals emergent cell patterning and tissue shaping. A,B,D) initial AS cells geometry with cells coloured according to their arrest time as parameters are varied: A, varying cMax and cFthr; B, varying alpha and beta; E, varying epidermis width and ES stiffness. Colour scale minimum and maximum are set for each simulation as the earliest and latest arresting cell time of that simulation. Cells in white do not arrest within the simulated time. C,D) Snapshots over time with larger epidermis along the AP (C) and DV (D) axis. E) Snapshots over time with larger epidermis along the DV axis and a computer-generated AS tissue discretisation in which AS cells are all almost equal to each other.

Next, we tested whether and how ANPi and ANPa production rates affected the patterned arrest of AS cell pulsing. To facilitate this analysis, we sped up simulated closure progression by fixing cMax = 2 and cFthr = 1.5. Under these conditions, the sequential nature of pulsation arrest turned out to be remarkably robust towards changes in ANPi and ANPa production rates (Fig. 5B). Obviously, oscillation arrest could only be monitored for parameter combinations that did lead to oscillations in first place, such as low ANPi production. In this case, the opening in addition did not contract. Similarly, high ANPi production rates caused absence of oscillations, but for another reason: Many cells arrested oscillating before ES tissue relaxation began. In contrast to cases with low ANPi production, this situation produced a strong tissue contraction,

Finally, we explored to what extent the mechanical properties of the ES tissue would affect the oscillatory behaviour and the pattern of contractility arrest of AS cells. Therefore, we fixed the actomyosin and mechanical parameters defining the AS tissue and selectively varied the resistance of the elastic ES tissue by modulating its geometry and its material parameters. First, we reduced the initial resistance by simply enlarging the simulated ES tissue (Figure 5C-D, Movies 6 and 7). As expected, this caused increased AS-shrinkage already before relaxation began (Fig. 5C, first two time points). If enlargement and thus resistance reduction was only implemented along one axis, the AS tissue shrank faster along this axis which altered AS tissue shape accordingly (Fig. 5C-D). Notably, not only the shape of the closing AS tissue varied, but also the pattern of sequential AS cell pulsation arrest (Fig. 5C-D, third and fourth timepoints). To further explore this effect of the epidermis tissue on AS cell actomyosin dynamics we systematically varied the ES tissue width along the DV-axis and the Young’s modulus of the ES tissue (Fig. 5E). So far, we had been using an ES tissue stiffness of E=1kPa. If the tissue was stiffened by increasing E, closure eventually could not proceed far enough for pulsation arrest to occur. Conversely, if the tissue was softened by increasing its width, closure eventually proceeded so fast that cells became fully covered by F-actin without ever oscillating. When the tissue was simultaneously hardened and enlarged, cells at the tissue boundary oscillated again and regained sequential arrest. Not surprisingly for a linear material, the effects of stiffness and width equilibrated. Intriguingly however, in case of low values for E and width, the direction of pulsation arrest progression turned by 90 degrees within the tissue as compared to the pattern emerging with high values for stiffness and width. Taken together, these data show that the mechanical properties of the ES tissue fundamentally influence the AS tissue shape and its patterning with respect to the oscillatory behaviour of individual AS cells in space and over time.

To estimate the extent to which our results were specific to the particular *in vivo* geometry we had selected, we repeated critical simulations on artificial geometries, where all AS cells were regularly arranged and identical in shape. We found the same effects on AS tissue shape and patterning but with the resulting patterns being much more regular than with the *in vivo* geometries (Fig 5F; Appendix Supplementary Methods).

### Simulations reproduce *in vivo* mutant AS cell behaviour

Starting from a scenario best reflecting wild type AS cell dynamics (Fig. 2D, Movie 3, Fig. 6B), and then varying the model parameters, our simulations produced a range of interesting cellular behaviours. We therefore checked if any of these simulations would reproduce mutant phenotypes as observed *in vivo. In vivo*, MyoII activity levels in AS cells have been manipulated by genetic interference or by interference with the MyoII phosphorylation levels either via ectopic expression of the inhibiting MyoII phosphatase (MbsN300) or the activating MyoII light chain kinase (ctMLCK) (Franke *et al*, 2005; Fischer *et al*, 2014; Saias *et al*, 2015). MbsN300 expression produced larger AS cells and tissue softening. Subsequently, a highly irregular pattern of apical AS cell constriction emerged. Conversely, ectopic ctMLCK expression produced smaller AS cells with longer-lasting contractile events and higher tissue tension. In our simulations, varying the threshold F-actin activity for contractility (cFthr) and the maximal contractility (cMax) mimicked loss and gain of MyoII activity. Activity gain correlated with increased cMax and decreased cFthr. This produced a phenotype very similar to that of ectopic ctMLCK expression: AS cells were smaller and exhibited longer-lasting and stronger contractile events (Fig. 6C, Movie 8). To mimic MyoII loss of activity, we reduced cMax. This produced a phenotype similar to that of ectopic MbsN300 expression, leading to AS cells that hardly contracted (Fig. 6D, Movie 9). This phenotype resembled another *in vivo* mutant phenotype that was observed in embryos maternally mutant for *rhoGEF2*, encoding the *Drosophila* Guanine nucleotide exchange factor of Rho GTPases (Azevedo *et al*, 2011). In these mutants, actomyosin coalescence significantly decreased and hardly any contraction occurred. Our simulations also phenocopied these effects. This is fully consistent with the suggested role of RhoGEF2 in promoting the association of MyoII with F-actin (Padash Barmchi *et al*, 2005). In turn, RhoGEF2 over-expression *in vivo* caused increased actomyosin coalescence in AS cells and led to contractile events that lasted longer and occupied larger subcellular regions, when compared both to wild type and to ctMLCK expression. Our simulations showed similar behaviour when we mimicked RhoGEF2 overexpression by increasing cMax (Fig. 6E, Movie 10). Thus, they also provide an interpretation for the DRhoGEF2 over-expression phenotypes indicating that increased MyoII recruitment is sufficient to reproduce the phenotype.

**Figure 6:**
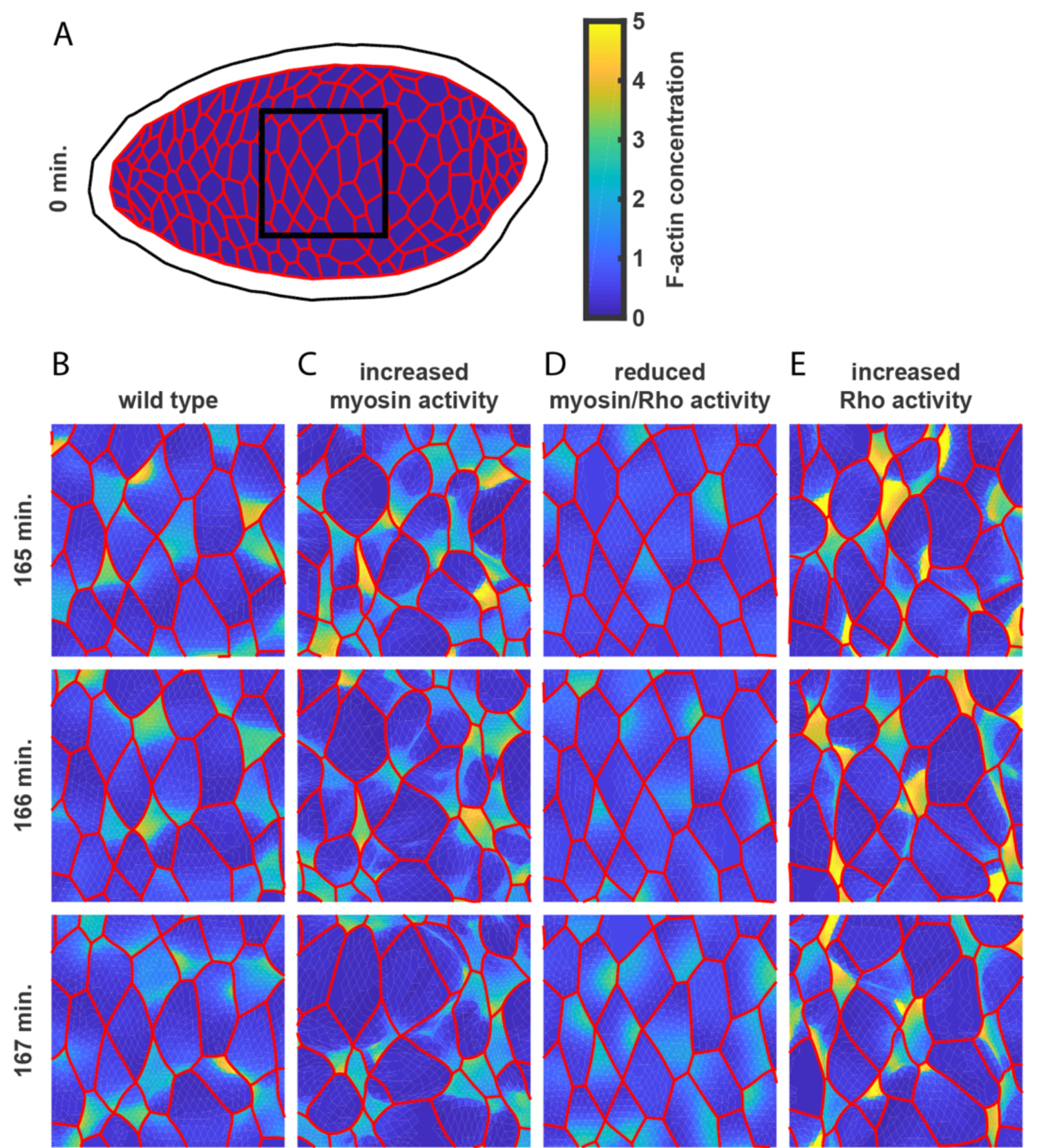
Simulations phenocopy wild type and mutant scenarios. A) initial geometry, cropped region (black square) and colour-scale of all subsequent time snapshots, chosen to show oscillatory dynamics. Simulations correspond to: B) wild type (same parameters as Fig. 2C,D); C) embryos with expression of ctMLCK (same parameters as B, except cFthr = 1, cMax = 2.5); D) embryos with expression of MbsN300 or maternal mutant for DRhoGEF2 (same parameters as B, except cMax = 0.2); E) embryos with DRhoGEF2 over-expression (same parameters as B, except cMax = 2).

In summary, our simulations indicate that varying actomyosin parameters at the subcellular scale is sufficient for the emergence of mutant phenotypes observed *in vivo* and they thus provide a clear interpretation.

## Discussion

Our FE modelling approach enabled a systematic multiscale exploration of the mechanism integrating repetitive apical AS cell surface constrictions occurring in the minutes range with simultaneous gradual apical constriction occurring over a time period of 3-4 hours to drive DC. The approach combines basic sub-cellular biochemistry with mechanical cell- and tissue behaviour. Our simple model predicts known behaviours to be emergent rather than the result of specific regulation by biochemical signalling. Biochemistry modelling is based on our experimental results, showing that F-actin oscillations are similar in presence and absence of MyoII activity. This differs from the mechanisms proposed for other tissue development processes employing oscillating cells for morphogenesis, such as the extending germ band (Munjal *et al*, 2015). There, mutant analysis suggested that oscillatory actomyosin dynamics depend on positive and negative biochemical feedback between MyoII advection and dissociation rates. Our model provides an alternative interpretation to consider, showing that under certain F-actin nucleation conditions, MyoII activity generates oscillatory F-actin dynamics without the need for biochemical feedback. Thereby, the contractility state of a given cell, indirectly influences the F-actin dynamics of neighbouring cells by changing their geometry. In other words, dynamic geometry can substitute biochemical signalling. In view of the predictions of our model, it would be interesting to see if during germ band extension, the absence of oscillatory dynamics in cells lacking MyoII activity can be compensated by altering F-actin production rates.

Currently our model is lacking the actomyosin ring that *in vivo* starts forming at the AS cell-cell junctions during DC onset and becomes increasingly prominent as DC progresses (Blanchard *et al*, 2010). In germ band extension, a non-uniform distribution of this actomyosin pool guides directional contractility (Rauzi *et al*, 2010). Its uniform distribution in AS cells could serve as a clutch that together with the medial-apical actomyosin pool generates a ratchet system or simply provide an additional, additive force. Arguing against the former, this pool is absent at the early phases of DC and only gradually builds up during the process (Blanchard *et al*, 2010). Furthermore, a recent publication linked this actomyosin activity to the control of junctional length and integrity (Sumi *et al*, 2018). Consistently, our model reproduces all *in vivo* behaviours found at the sub-cellular, cellular and tissue scale without this actomyosin pool. Even more, our model reproduces much of the mutant phenotypes observed *in vivo* by simply varying the parameters reflecting the affected biochemical *in vivo* activity. Thereby, the model not only shows sufficiency but suggests causative mechanisms for phenotype development, defining a mechano-chemical parameter space within which the cells need to reside for a given phenotype to occur. For example, previous *in vivo* modulation of MyoII activity produced phenotypes that could not be explained in a simple way and were hypothesised to be caused by pleiotropic effects of MyoII activity on cellular function (Fischer *et al*, 2014). Our model reproduces these phenotypes without addition of further complexity. Of course, the predictions of the model now ask for more comprehensive analysis of the *in vivo* systems. To address the contributions of the junctional and the medial-apical actomyosin pools and their mechanical and chemical parameters to DC *in vivo*, will require spatially and temporally controlled, acute protein interference as well as careful biophysical measurements for example of tissue stiffness over time. Such analysis may well lead to a reinterpretation of some of the cellular phenotypes and the associated closure defects.

Currently, the initial event triggering DC is not known. Regulation of the pulsed contractile forces of AS cells can be excluded, as this critical activity starts prior to dorsal closure initiation. Some other change thus must occur in the AS or the surrounding ES tissue. In our model, a gradual relaxation of the ES tissue was sufficient to allow DC initiation and propagation. Such a scenario was previously proposed based on the observation that during the process the ES cells gradually elongated in the direction of tissue closure (Riesgo-Escovar & Hafen, 1997). Such tissue relaxation is generally presumed to be a fundamental mechanism of embryonic tissue morphogenesis and wound healing (Razzell *et al*, 2014; Bambardekar *et al*, 2015; West *et al*, 2017). Consistently, DC arrests if ES cell elongation is prevented by prohibiting Dpp secretion from the DME cells or by mutating the Rho/Rac effector target Pkn (Riesgo-Escovar & Hafen, 1997; Lu & Settleman, 1999). The importance of this mechanism was questioned as the selective rescue of global Dpp interference implied that Dpp signalling in addition regulates AS cell constriction, which may well be the critical activity triggering DC (Garcia Fernandez *et al*, 2007; Zahedi *et al*, 2008). However, our direct, AS tissue-specific interference with Dpp signalling, contradicts these findings, bringing Dpp-mediated ES tissue relaxation as DC initiator back into focus.

This finding in addition supports a further prediction of our simulations, namely that the sequential arrest of AS cell pulsation described *in vivo*, is not brought about by a Dpp gradient emanating from the DMEs, but simply is an emergent property of the tissue systems mechanics (Solon *et al*, 2009; Blanchard *et al*, 2010). Our simulations could reproduce the in vivo pattern of pulsation arrest with AS cells at the ES/AS tissue boundary arresting first and cells in the tissue centre last. Thereby, AS cells arrested in a contracted state, with their entire surface covered by F-Actin as previously described (Blanchard *et al*, 2010; Jayasinghe *et al*, 2013; Machado *et al*, 2015). *In vivo*, this effect was assigned to differential biochemical regulation over time of one or more factors of the actomyosin system. In our model, the behaviour emerges without any specific regulation. The shrinking AS cells eventually reached sufficient ANPa-levels for F-actin structures to cover their entire surfaces. This effect relied on our assumption that the amount of ANPi remains constant as the cell surface area gradually shrinks. *In vivo*, this would mean that the amount of F-actin nucleator reaching the apical plasma membrane is constant over time, a reasonable scenario, assuming directed transport or tethering of rate limiting factors. It is plausible, that the sequential nature of this effect in our simulations starts emerging at DC onset, due to the outermost AS cells on one side being connected to passive ES cells. This means that after each contraction event, these boundary cells have fewer actively contracting neighbours than AS cells inside the tissue that drive their subsequent expansion. Consequently, they reach a fully contracted state much earlier. This effect propagates as in the contracted state the boundary cells in turn have become passive and no longer contribute to the stretching of the next inner AS cells. This generates a new, mechanical boundary that similarly accelerates pulsation arrest of the next inner AS cells. In this way pulsation arrest propagates further into the tissue over time.

Previously, we had discussed that this arrest in a contracted state could be compensating the surface tension-loss of the shrinking tissue (Solon *et al*, 2009). This is consistent with a subsequently proposed mechanism, where contracting cells stretch neighbouring cells to open mechanically gated ion channels, which induces these cells to contract in turn (Hunter *et al*, 2014). However, we show that F-actin oscillation dynamics are not affected in tissues lacking mechanical input due to MyoII activity depletion. The reported Ion channels must be serving some other important function. This is consistent with another issue addressed with our computational modelling of DC - the coordination of neighbouring AS cell surface oscillations. Quantification of experimental data indicated that neighbouring cells generally oscillate in anti-phase (Solon *et al*, 2009), or form rows of cells that contract at the same time (Blanchard *et al*, 2010). In our simulations various coordination patterns emerged depending on the biochemical and mechanical parameters in the complete absence of any biochemical coupling between neighbouring cells.

Intriguingly, in a given AS tissue with a fixed set of biochemical and mechanical parameters the pattern of the sequential pulsation arrest can differ fundamentally, solely depending on the local elastic properties of the surrounding ES tissue. Simultaneously, these properties define AS tissue shape. It is thus possible that also *in vivo* changing mechanics of surrounding tissues critically affect the patterning and morphogenesis of a tissue without it changing its molecular state. Hence, when exploring tissue and organ morphogenesis one cannot avoid considering cooperative effects of simultaneously developing tissues. This point is interesting also from an evolutionary perspective suggesting that morphological changes of a tissue are not necessarily driven by tissue-autonomous changes in gene expression.

Altogether, our simple model predicts a range of known behaviours to be emergent, which indicates the predictive power of our simulations. Importantly, these predictions emphasise the importance of identifying *in vivo* the contribution of geometry and of cell- and tissue scale mechanics to sub-cellular biochemical dynamics such as F-actin pattern modulation. In principle, our model can be adjusted to simulate other oscillatory tissues undergoing morphogenesis. It would for example be interesting to explore whether and to what extent it could reproduce gastrulation and germ band extension, the other classical morphogenetic processes that have been intensively studied in flies.

While our results show the immense potential of FE-based modelling in enabling more complex computational exploration of living systems, they also stress the need for a holistic analysis at single cell and tissue scales of entire tissue communities for a more profound understanding of morphogenesis.

## Materials and Methods

### Fly strains

Fly strains used in this study are listed in Supplementary Table 1.

For AS tissue-specific interference with Dpp signalling, 332.3Gal4 (Wodarz *et al*, 1995) was used to drive tkv^act^ expression and P0172Gal4 was used to drive UAS-Dad (Tsuneizumi *et al*, 1997). Notably, expressing UAS-Dad via the 7KGal4, KrüppelGal4 (Castelli-Gair *et al*, 1994) or the combined dpp[4B] and dpp[8B]Gal4 drivers, also did not affect DC. This was not due to a lack of functionality as this combination caused adult fly lethality (data not shown).

### Confocal microscopy and image processing

Embryos were prepared for imaging as previously described (Jankovics & Brunner, 2006). Imaging of all embryos was performed at 23-25°C using spinning-disk confocal microscopes (Zeiss Axio Observer.Z1 or custom-modified Leica DM IRBE, equipped with iXon3/888 and Neo sCMOS cameras and controlled by ANDOR IQ software or VisiScope Confocal-FRAP Cell Explorer). Objectives used: 25X (for dorsal closure overview), 40X, 63X and 100X. Z-planes were acquired every 2µm and maximum-intensity z projections were analysed. For higher resolution of LE cells, amnioserosa cell and MyosinII dynamics (63X and 100X objectives) z-planes were acquired every 0.5-1µm, and single planes or maximum-intensity z projections of relevant planes were analysed.

Image processing and maximum intensity z-projections were done using ImageJ or MATLAB (MathWorks).

### F-actin dynamics analysis

F-actin dynamics were quantified using maximum intensity z-projections of wild type and AS-SqhKO movies of AS tissue. Using ImageJ, we first measured mCherry-moesin fluorescence intensities in ROIs within AS cells every 30s throughout the movie sequences. The intensity oscillations were then “detrended” by subtracting the mean intensity value individually for every intensity profile. To determine the periodicity of F-actin nucleation bursts, we divided the movie frames into time windows of 900s that were shifted by 30s. For every 900s window from intensity profile auto-correlation was performed on the intensity profile (autocorr function in MATLAB). The average distance between the peaks in the auto-correlation function was taken as a period for that time window. Peaks were found using the “findpeaks” function in MATLAB without any threshold for peak heights. We then plotted period distributions according to their relative frequency (Fig. 1B).

Wild type: n=13508 CCFs, 155 unique cell pairs (= 310 cells), 3 embryos; AS-SqhKO: n=22712 CCFs, 205 unique cell pairs, 5 embryos

## Acknowledgements

We thank Tinri Aegerter and Darren Gilmour for critical reading of the manuscript. This work was supported by a SystemsX.ch grant to DB and CA as part of the MorphogenetiX project and by the Bundesministerium für Bildung und Forschung 031A494 to RSS.

The authors declare that they have no conflict of interest.

